# DNA methylation mediates transcriptional stability and transposon-driven *trans*-regulation under drought in wheat

**DOI:** 10.64898/2025.12.04.692301

**Authors:** Isaac J. Reynolds, Liam J. Barratt, Andrea L. Harper

## Abstract

Bread wheat (*Triticum aestivum*) is a major component of half the global population’s diet, but increasingly frequent droughts threaten its productivity and food security. While massive transcriptional reprogramming under drought in wheat seedlings is well characterised, DNA methylation’s contribution remains poorly understood. Using paired whole-genome bisulphite sequencing (WGBS) and RNA-seq before and after drought stress in wheat landraces, we probed the nuanced role of DNA methylation in the drought response, uncovering antagonistic trends between cytosine contexts and novel mechanisms, with the ROS1a family potentially playing a key demethylation role under drought. Examination of gene methylation profiles revealed that gene body methylation was strongly positively correlated with gene expression but negatively with stress responsiveness, simultaneously identifying that gene body differentially methylated regions (DMRs) targeted stress-associated genes. Many DMR-associated genes maintained consistent transcription under stress, suggesting a stabilising role for DNA methylation. Most DMRs localised to intergenic regions and transposable elements (TEs), with the ancient LTR retrotransposon *RLX_famc9* emerging as a critical target of differential methylation under drought. We propose a model in which the *RLX_famc9* family, enriched in differential methylation and exhibiting substantial sequence similarity to drought-responsive genes, is involved in the *trans*-regulation of stress-associated genes under control conditions through the generation of regulatory siRNA precursors, a mechanism suppressed by drought-inducible hypermethylation. Our findings suggest an intricate regulatory role of DNA methylation under drought, with genic DNA methylation promoting high, stable expression, ROS1a glycosylases coordinating targeted demethylation, and methylation-controlled TEs modulating the expression of downstream genes in *trans*.

## Introduction

As the climate changes in lockstep with growing populations, resource scarcity and increasing aridity of arable land threaten global agriculture’s ability to meet rising nutritional demand. Crops are continually exposed to a variety of stresses in the environment: mitigation is paramount for obtaining sufficient yield and ensuring food security. Environmental challenges like intense heat, salinity, and drought trigger dynamic molecular changes in plants as they adapt to survive and reproduce. Environmental cues and stress responses in plants prompt transcriptional reprogramming steered by regulatory genes and epigenetic mechanisms ^1–4^. Epigenetic changes – including alterations to chromatin accessibility, production of small RNAs (sRNAs), or RNA and DNA methylation – are known to cause shifts in gene expression ^5,6^, though the extent and effect of these shifts remains a subject of debate. The plant epigenome may give rise to greater phenotypic plasticity, enabling plants to adopt more beneficial phenotypes under stress and ultimately attain greater survival and yield in adverse conditions ^7,8^.

Bread wheat (*Triticum aestivum*) is a staple grain crop relied upon by billions globally as a major source of calories, protein, and fibre ^9,10^; further crop development to ensure its yields among increasingly harsh climates is fundamental to achieving and maintaining food security across the world. In order to meet the nutritional demand of the future, predictions suggest that the yields of crops like wheat need to increase by up to 50% ^11,12^, though some estimates have suggested that even this will not be sufficient or sustainable ^13^, prompting the need for new avenues of crop improvement.

Among the most critical threats to wheat production in much of the world is climate-change-driven drought, with water deficits shown to reduce grain and grain protein yields by almost half ^14,15^. Given wheat’s importance to much of the world’s dietary intake, sustained reductions in yield would have widespread consequences for food security, agriculture, and economic stability. In much of the world, major crops including wheat are grown under rainfed conditions ^16–18^, leaving them highly vulnerable to unexpected shifts in precipitation, even where supplemental irrigation is available. In addition, early drought stress continues to be an area of increasingly intense research, with a number of studies in recent years exploring the impacts of early drought on wheat and stressing the importance of identifying tolerant varieties ^19–21^. However, the higher regulatory molecular basis of their ability to respond to drought remains underexplored. Periods of water shortage are becoming increasingly common throughout the year ^22^, overlapping with the early stages of spring wheat growth: following current trends, this will continue to intensify as climate change progresses, and so understanding the mechanisms underlying plant responses to drought at the genomic level will enable us to better exploit their potential and bolster their survival.

Genome-wide epigenetic marks (e.g. DNA methylation, histone modification, small RNAs, chromatin remodelling) are associated with almost every aspect of environmental response and plant development ^23,24^. DNA methylation, where a methyl group is added to the fifth position of cytosine residue (5mC) is known to play a role in the regulation of gene expression and so is likely involved in abiotic stress responses. In plants this is mediated by the RNA-directed DNA methylation (RdDM) and demethylation pathways through interactions between small RNAs, protein complexes, and DNA methyltransferases or glycosylases. Cytosine-based DNA methylation can occur at three cytosine contexts: CG, CHG, or CHH (where H represents any base other than G). DNA methylation likely has different impacts on transcription depending on its localisation; methylation markers in the gene body may be positively associated with expression ^25^, while methylation at promoter regions and beyond is typically associated with gene and transposable element silencing ^24^.

Investigation into wheat’s methylome and its role in stress responses is limited; largely, current research has relied on techniques like methylation sensitive amplification polymorphism (MSAP) to profile the methylome and identify changes at broad regional levels ^26^, have targeted specific genes ^27^, or used lower-resolution approaches like meDIP-Seq. Genome-wide approaches to studying the wheat methylome have been used in studying the biotic stress response ^28^, but the use of techniques like whole-genome bisulphite sequencing (WGBS) or long-read sequencing remain rare in wheat, in part due to the extensive cost and time of sequencing and analysis for the large genomes of complex polyploids. However, genome-wide approaches have been used to investigate the drought response methylome of crops like rice ^29^ and maize ^30^, identifying correlations with gene expression and differentially methylated genes that may be involved in the drought response.

Despite making up a large proportion of plant genomes and being associated with a greater proportion of differential methylation, the regulatory potential of transposable elements (TEs) remains underexplored. Most TEs in plant genomes are highly and stably methylated, but some have been identified as being involved in stress responses through *cis*- or *trans*-regulation: the *Helitron* TE family *ATREP2* is likely integral for induced resistance to herbivory in *Arabidopsis thaliana* ^31^, while non-coding RNAs derived from TEs have been identified as important for abiotic stresses in Moso bamboo ^32^ and maize ^33^.

The present work represents a novel integration of transcriptomics and whole genome bisulphite sequencing data to explore the thus-far understudied role of epigenetic regulation in wheat’s drought response, with a focus on changes at genes and transposable elements. By examining changes in methylation dynamics across the genome and identifying correlations with transcriptional shifts, we aimed to elucidate this regulatory layer of abiotic stress responses and identify potential avenues for crop improvement under challenging environmental conditions.

## Results

### Phenotype

Phenotype data involving the plants used in this study was previously reported ^21^; drought treatment was found to significantly negatively impact fresh (*p* = 1.01e-09) and dry weight (*p* = 7.75e-06) when comparing the treatment and control groups (one-way ANOVA). Accessions with high mean biomass retention (>70%) in the drought group compared to their controls were used for WGBS. Samples were taken from plants at treatment onset (BD; Zadok’s GS13) and 10 days post treatment (AD).

### Transcriptional reprogramming in spring wheat seedlings under drought

This present study used 328.1 Gb of raw data generated using the Illumina paired-end Novaseq 6000 platform as per *Barratt & Reynolds et. al 2023*. 7.264 x 10^8^ raw reads across 14 samples were used with an average of 97.3%/ 92.6% bases having a q-value of ≥20/ ≥30 respectively and an error probability of 0.03. Mean GC content ranged from 53-57%. Raw reads were assessed using FastQC and pre-processed using Trim-Galore.

Counts from the 14 samples (7 per group) were transformed using the vst() function of DESeq2 for exploration using principal component analysis (**Figure 1A**): PC1 (64% variance) clearly distinguished between samples taken before (BD) and after drought (AD).

**Figure 1:**
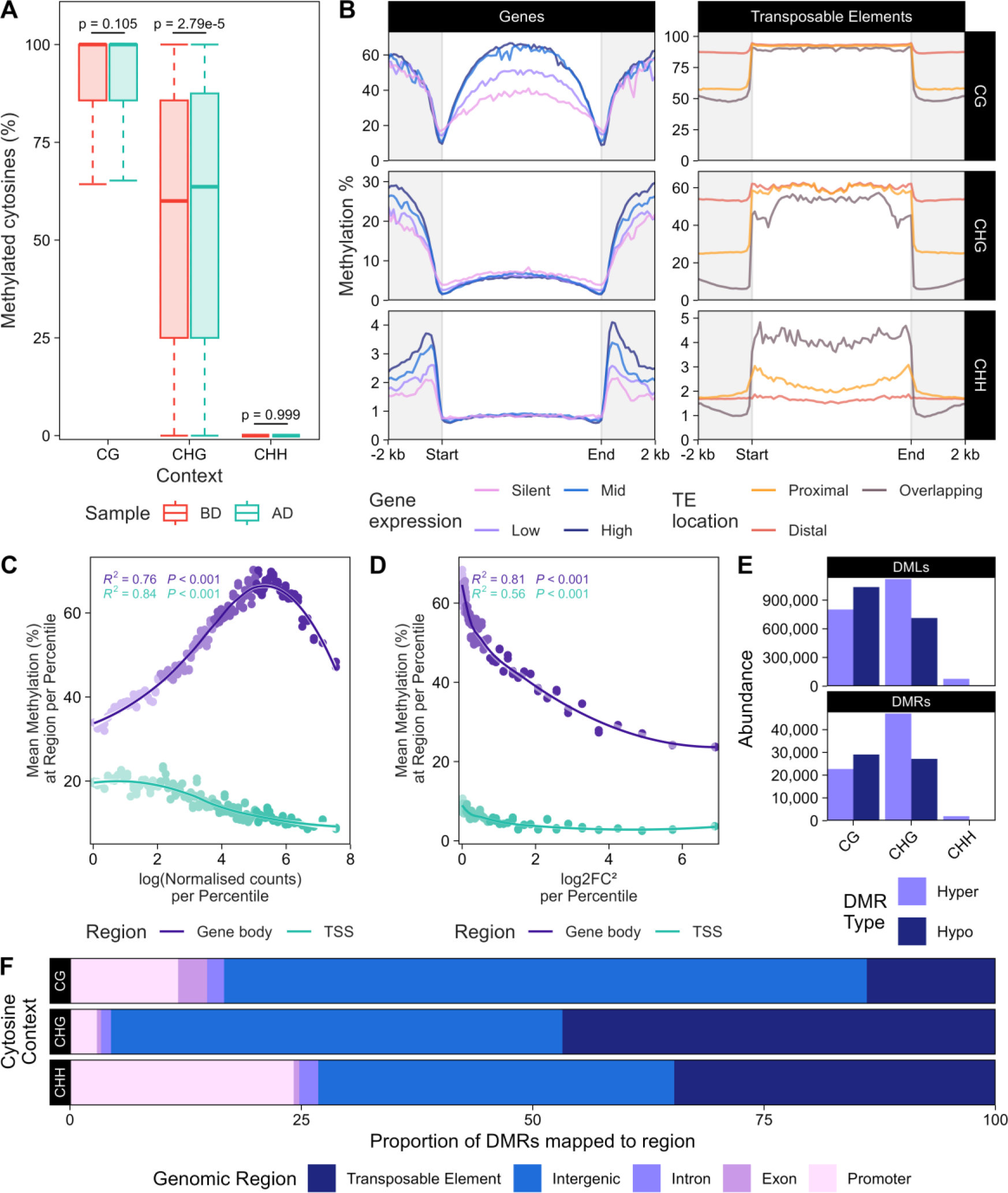
**(A)** PCA of global gene expression patterns identified that PC1 was the main driver in the dataset and strongly distinguished between samples before and after drought treatment. **(B)** GO enrichment was performed on both upregulated and downregulated gene sets using ClusterProfiler. A selection of stress-associated GO terms is provided. Complete GO enrichment results for differentially expressed genes can be found in **Supplemental Data S3. (C)** Gene expression of major wheat methyltransferase and DNA glycosylase families was compared before and after drought. *** represents p < 0.001; NS represents no significant difference. **(D)** ROS1a TILLING lines were used to examine the role of ROS1a-mediated demethylation in determining drought tolerance. WT and MUT refer to wild-type and mutant BC_1_S_1_ lines. Dashed line represents the average nDW of Cadenza under the same conditions for comparison.

Differential expression analysis was carried out to identify genes that responded to drought at the seedling stage, identifying 13,932 DEGs. 58% were upregulated after drought. Stress response GO terms were enriched in upregulated genes (**Figure 1B**) with ‘response to water deprivation’ (FDR-adjusted *p* value = 1.77e-66) the most significantly enriched, accompanied by ‘leaf senescence’ and ‘stomatal movement’. GO terms associated with processes like ‘photosynthesis, light reaction’ (*p* = 2.69e-56) and ‘water transport’ (*p* = 2.16e-08) were enriched in downregulated genes. GO enrichment results for upregulated and downregulated gene sets are available in **Supplemental Data S2**.

DNA methyltransferase and glycosylase gene family expression was compared at the family level before and after drought (**Figure 1C**). CMT2 expression was significantly higher (*p* = 1.81e-05; *Wilcoxon ranked-sum test*) after drought treatment, while all other families showed increased but non-significant changes. ROS1a expression was significantly higher following drought (*p* = 2.99e-04; *Wilcoxon ranked-sum test*); though the three other DNA glycosylase families fluctuated in expression, none were significant.

### ROS1a knockouts may have homoeologue-specific effects on drought tolerance

Mutant lines with single ROS1a homoeologue knockouts (ROS1a-5A^mut^, ROS1a-5B^mut^, and ROS1a-5D^mut^) were used to explore the role of ROS1a in the drought response (**Figure 1D**). ROS1a^mut^ displayed contrasting trends when compared to both wild-type Cadenza and homozygous ROS1a^WT^ lines with the same mutation load.

Almost all lines exhibited higher drought tolerance than Cadenza. Mutant lines generally exhibited nDW reduction compared to WT. Genotype had a significant effect on normalised dry weight (ANOVA*: p =* 0.0139) in ROS1a-5A plants; ROS1a-5A^mut^ displayed significantly higher nDW than ROS1a-5A^WT^ in post-hoc Tukey testing (+9.1%, *p* = 0.0129).

Among ROS1a-5B plants, genotype also had a significant effect on nDW (ANOVA*: p =* 0.0118); ROS1a-5B^WT^ nDW was significantly higher than Cadenza (post-hoc Tukey test; *p* = 0.0124). ROS1a-5B^WT^ plants displayed a 32.4% higher mean nDW than ROS1a-5B^mut^ plants and 49.7% higher than Cadenza.

Similarly, genotype had a significant effect on nDW in ROS1a-5D plants (ANOVA*: p =* 0.0442), with Tukey tests identifying significant differences between ROS1a-5D^WT^ and Cadenza (*p* = 0.0462). ROS1a-5D^WT^ plants displayed a mean 10.2% increase in nDW compared to ROS1a-5D^mut^ plants.

Examination of all lines together identified a significant effect of genotype on nDW (Aligned Rank Transformed (ART) ANOVA*: p =* 0.000196), alongside a significant interaction term (*genotype x homoeologue*) (ART ANOVA*: p =* 0.000226). Post-hoc testing identified that Cadenza was significantly different to both WT (*p* = 0.000650) and mutant ROS1a plants (*p* = 0.000665), but ROS1a WT and MUT plants were not significantly different (*p* = 0.889) to eachother. A slight decrease (1.7%) in nDW was observed in ROS1a MUT lines compared to their WT siblings.

### BS-Seq and differential methylation of the wheat genome under drought

Before drought (BD) and after drought (AD) samples were sequenced across 5 lanes, generating 799.8 Gb of raw data, consisting of ∼5.3×10^9^ reads.; an average of 94.7%/ 87.4% of bases had q-values of >20/ >30 and an error probability of 0.03. GC content ranged between 26.03% - 26.64%.

Average unique mapping rates to the RefSeq IWGSC v1.1 assembly (International Wheat Genome Sequencing Consortium; http://wheatgenome.org) were 72.9% (BD) and 72.0% (AD). Remaining reads were discarded due to alignment score or multi-mapping. Average coverage was 13.6X (BD) and 14.5X (AD). Proportions of methylated cytosines for each context (CG, CHG, and CHH) were 87.0%, 53.8%, and 1.6% and 87.1%, 54.8%, and 1.7% in BD and AD respectively. Methylation calls at sites with fewer than 3 reads were removed from further analysis.

Cytosine contexts displayed distinct methylation trends. Statistically significant differences between BD and AD samples were only observed between CHG sites (+1.2%, Tukey*: p =* 3.20e-07) using an ANOVA (**Figure 2A**). Little preference was observed among the methylation state of context in its triplet base contexts; for example, average methylation at each CG triplet (e.g CGT, CGG, CGC etc.) ranged between 86.3 - 90.1%.

**Figure 2:**
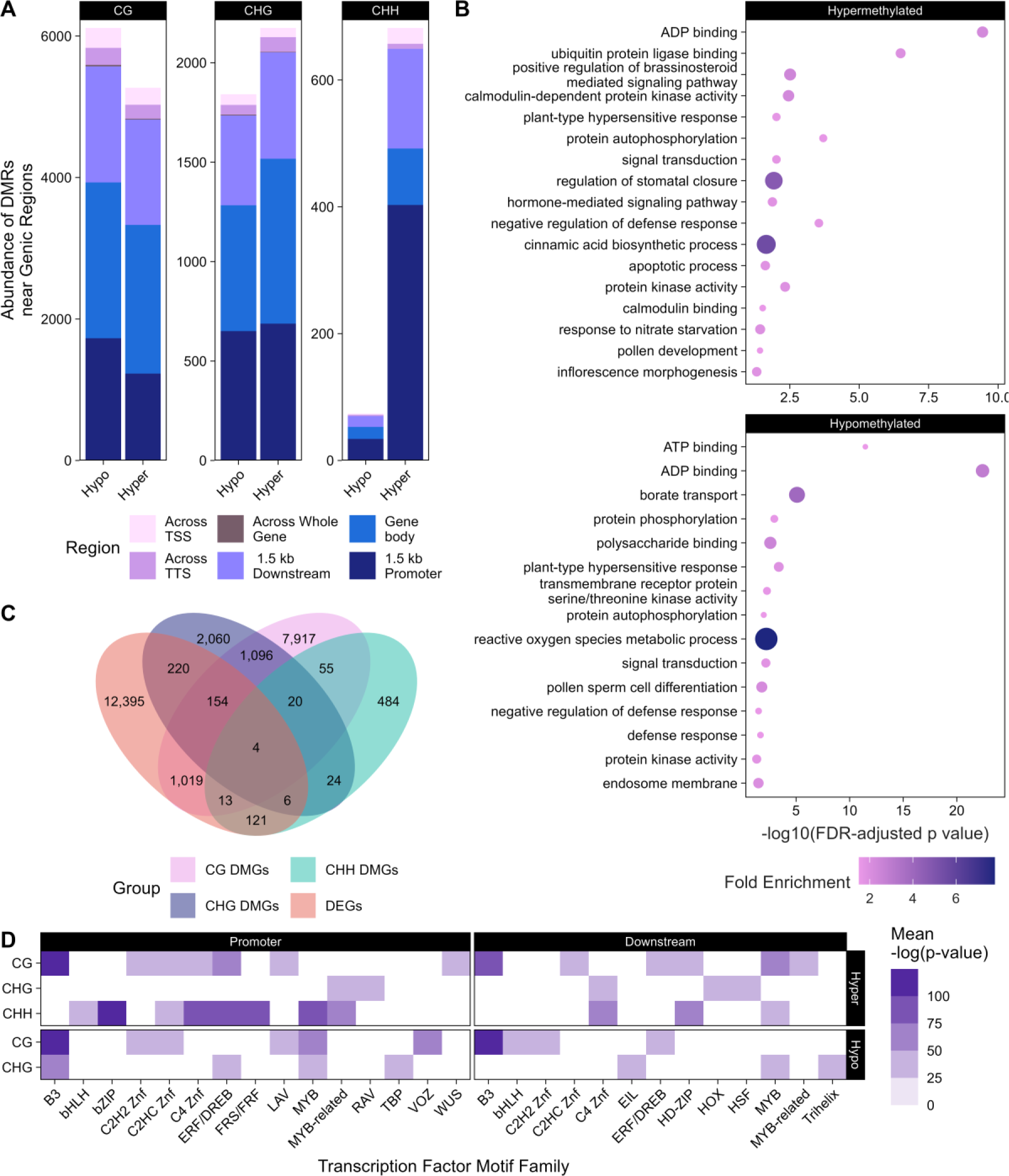
**(A)** A sample of ∼5% of methylation calls were sampled from each file and visualised using boxplots. Methylation at each context ranged from 0-100%. **(B)** Mean methylation across genes and transposable elements was calculated per 100 bins (20 bins per 2 kb flanking region, 60 bins per body). Genes were grouped into four expression quartiles and transposable elements were grouped by gene proximity (“overlapping genes”; <10 kb: “proximal”; >10 kb: “distal”). **(C)** Genes were grouped into percentiles by mean expression. Mean log(normalised counts) percentile was plotted against maximum CG gene body methylation (above) as well as average methylation at the TSS (below). **(D)** Genes were grouped into percentiles using absolute log₂FC (BD vs AD); the mean absolute log₂FC was plotted against the maximum mean CG gene body methylation (above) as well as average methylation at the TSS (below). (**E)** Differentially methylated loci (DMLs) and regions (DMRs) were summarised per context. (**F)** DMRs were mapped to genomic and genic features using the R package genomation.

Gene bodies displayed methylation depletion across the transcription start (TSS) or termination (TTS) sites, consistent increases towards the centre of the gene, and increasing methylation with distance from the gene body in the flanking regions (**Figure 2B**). For TEs, proximity to genes was a substantial factor in determining methylation profile across all three contexts; proximal and gene-overlapping TEs displaying substantially lower methylation levels in flanking regions than distal TEs.

Genes, grouped into percentiles by expression (**Figure 2B, 2C**) and absolute log₂Fold Change **(Figure 2D)**, were significantly associated with the peak of CG gene body methylation. Significant correlations were identified between the mean expression of percentile bins, their maximum mean gene body methylation (*p* < 0.001; R^2^ = 0.76), and the mean methylation at the bin containing the transcription start site (TSS) (*p* < 0.001; R^2^ = 0.84). Gene expression variability under drought by percentile was also significantly correlated with DNA methylation: average absolute log₂Fold Change of percentiles and their maximum mean gene body methylation (*p* < 0.001; R^2^ = 0.81) and average methylation at the TSS (*p* < 0.001; R^2^ = 0.56) both showed significant correlations. Gene body DNA methylation was positively correlated with expression, while TSS methylation was negatively correlated. High methylation at the TSS and gene body were weakly and strongly negatively correlated respectively with log₂Fold Change under drought. Only small variation was observed between BD and AD, though this may be an artifact of the quantile approach.

Differential methylation identified both differentially methylated loci (DMLs) and regions (DMRs). Raw differential methylation results are summarised in **Figure 2E**. CG DMRs (Δ Methylation threshold = 20%) displayed preferential hypomethylation (56.1% of 51,775 CG DMRs). Non-CG contexts yielded the most DMRs, with stronger bias towards hypermethylation; 62.9% of the 74,185 CHG DMRs (Δ Methylation threshold = 20%) identified were hypermethylated. CHH (Δ Methylation threshold = 10%) yielded the fewest DMRs (2,281), with 85.5% of these hypermethylated. CHG and CHH DMRs were on average 29.0% and 35.9% shorter than CG DMRs, while CHH DMRs were substantially more C-rich, containing over twice as many cytosines on average.

Mapping of DMRs to genomic features (**Figure 2F**) revealed that over 69%, 48%, and 38% of CG, CHG, and CHH DMRs were localised to intergenic regions. Gene-associated CHG DMRs most frequently localised to the gene body, while gene-associated CG and CHH DMRs primarily localised to the gene-flanking regions. Gene promoter regions were the dominant localisation of genic CHH DMRs (∼24% overall), and the second-most overlapped feature for genic CG (∼12%) and CHG (∼3%) DMRs. Transposable elements were the most frequent annotated regions for all contexts.

### Genic DNA methylation may play an influential but secondary role in the drought stress response

9.78% of DMRs were associated with promoter, exon, or intronic regions (**Figure 3A**). CG and CHG DMRs mapped in similar proportions to different genic regions, while CHH DMRs were dominant in promoter regions. Differentially methylated genes (DMGs) were defined as genes that overlapped with a DMR in the gene body or across its flanking regions.

**Figure 3:**
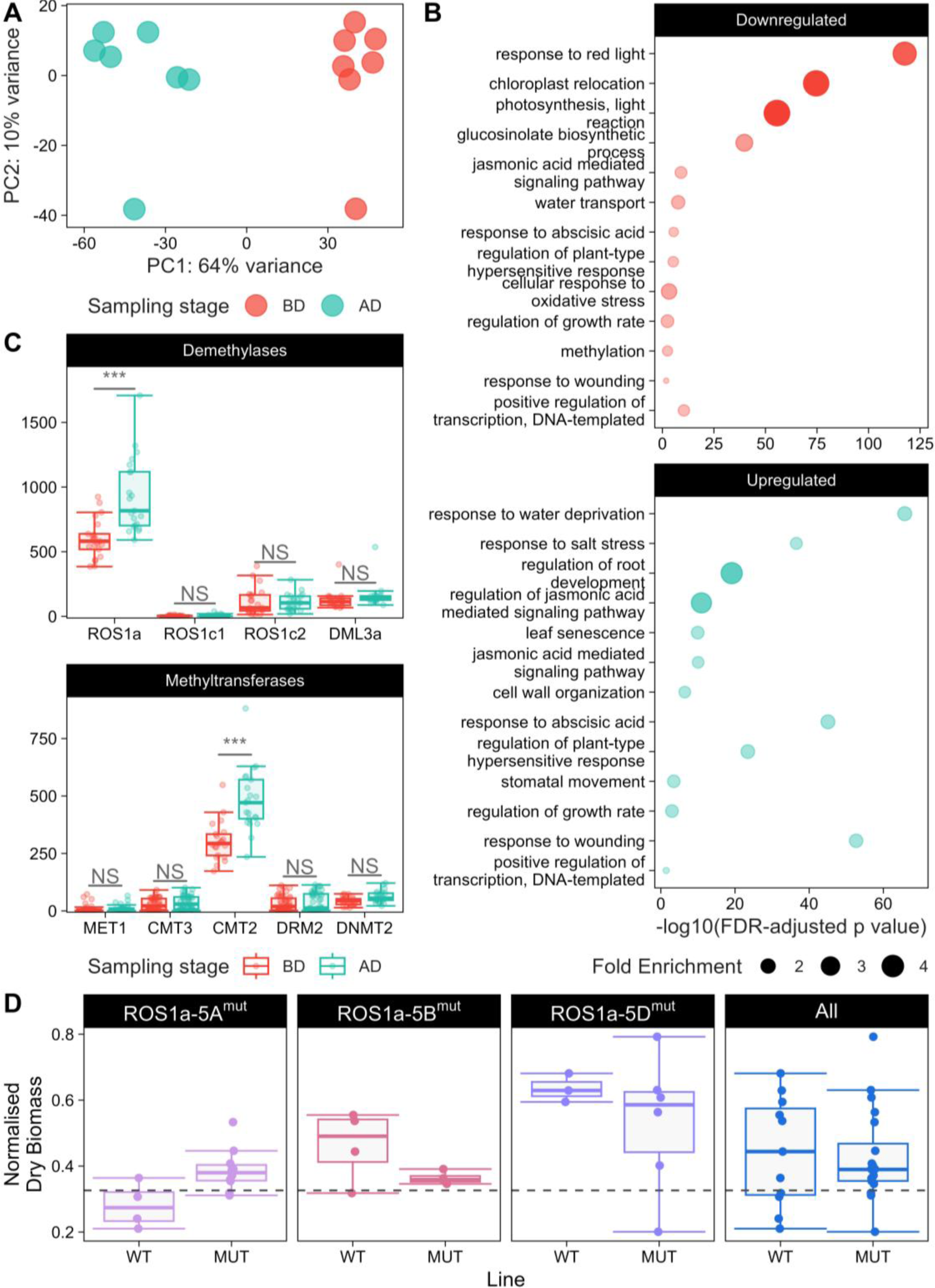
**(A)** DMRs mapped to genes and their flanking regions were broken down into locales based on their start and end coordinates relative to the gene annotation. **(B)** GO enrichment analysis was conducted using DMGs associated with gene body DMRs per type against a background of all genes with annotated GO terms. Size and colour represent fold enrichment. **(C)** DEGs and DMGs for all contexts were compared using a venn diagram. **(D)** Statistical motif enrichment of hypo- and hypermethylated DMR sequences in gene-flanking regions was carried out using HOMER. Results were summarised to the predicted family level. Darker cells indicate higher significance.

GO enrichment was performed using DMGs against all wheat genes annotated with GO terms (**Figure 3B**). Stress-associated GO terms were enriched in hypomethylated CG gene body DMGs, including terms such as ‘*reactive oxygen species metabolic process’* (GO:0072593), ‘*plant-type hypersensitive response’* (GO:0009626), ‘*hormone-mediated signalling pathway’* (GO:0009755), and ‘*regulation of stomatal closure’* (GO:0090333). 104 GO terms were enriched in hypermethylated CG gene body DMGs, including ‘*positive regulation of abscisic acid-activated signalling pathway’* (GO:0009789), ‘*response to salicylic acid’* (GO:0009751), and ‘*response to cold’* (GO:0009409). Both were enriched in ‘*protein phosphorylation’* (GO:0006468) and ‘*signal transduction’* (GO:0007165).

Enriched GO terms were also obtained among CHG DMGs: hypermethylated gene body DMGs were enriched in the terms ‘*potassium ion transmembrane transport’* (GO:0071805) and ‘*abscisic acid-activated signalling pathway’* (GO:0009738), while hypomethylated gene body DMGs were enriched in similar terms to their CG counterparts, including the terms ‘*signal transduction’* (GO:0007165) and ‘*plant-type hypersensitive response’* (GO:0009626).

No enriched terms were observed among CHH DMGs with gene body-associated DMRs. No enriched GO terms were observed among DMGs with DMRs in the flanking regions for any cytosine context.

### DEGs, DMGs, and DMRs are partially shared across contexts

DMGs of each context and DEGs substantially overlapped (**Figure 3C**). Four genes overlapped all: *TraesCS1D02G012000* (log₂FC = 5.65, FDR = 2.11e-10), a defensin-like AMP protein, *TraesCS2A02G313000* (log₂FC = -1.28, FDR = 1.1e-3), a cupin superfamily protein, *TraesCS5D02G317000* (log₂FC = -1.26, FDR = 0.038), a CRT/DRE binding factor protein, and *TraesCS7A02G566200* (log₂FC = 3.40, FDR = 4.1e-4), an ABC transporter G protein. DMG/ DEG overlaps displayed weak relationships between the expected effect of DMRs and log₂Fold Change (e.g. promoter hypermethylation repressing expression, or gene body hypermethylation enhancing it). Only 49.5% CG, 48.6% CHG, and 58.7% of CHH DMRs associated with promoters or gene bodies displayed the expected change in expression.

DMRs within 100bp of other DMRs, including across contexts, were largely positively correlated: of 10,108 CG-CHG proximal DMRs, 99.7% displayed similar methylation profiles, with 55.8% of these being hypomethylated. 88.48% of proximal CG-CHG DMRs were mapped to intergenic regions. Other comparisons yielded few proximal DMRs: only 57 CHG-CHH and 70 CG-CHH DMRs were observed, of which 79% and 9% displayed similar methylation profiles.

### Gene-flanking DMRs are enriched in transcription factor binding motifs

Overrepresented transcription factor binding motifs (TFBMs) were identified within DMRs localised to the flanking regions of genes for all three contexts. These regions were analysed with enriched motifs summarised to the family level. 43 unique significantly enriched motif families (*p* value ≤ 1e-10 & average log enrichment ≥ 2) were identified across all contexts (**Figure 3D**). Several TFBMs were associated with stress-associated families, including members of the *MYB, bHLH, ERF/DREB, bZIP*, and zinc-finger-containing families. The most significantly enriched family was *MADS*-box transcription factors (mean p-value = *1e-28*), driven by an abundance of *VRN1* and *REM* motifs. Several families were enriched in both upstream and downstream flanking regions as well as enriched in both hypo- and hypermethylated DMRs. Low abundance of hypomethylated CHH DMRs resulted in no enrichment of families.

### Differential methylation in non-genic regions

DMRs mainly localised to non-genic and unannotated regions; >40% overall mapped to TEs. 7,992 CG DMR-associated TEs (DMTEs) were identified, trending towards hypomethylation (54.6%). The RLG (Ty3-like, formerly known as *Gypsy*) superfamily was most abundant for CG among both methylation states, followed by RLC (Copia), and DTC (CACTA). More CHG DMRs (46.8%) were associated with TEs than CG DMRs (13.9%); of 47,014 CHG DMTEs, 67.8% were associated with hypermethylated DMRs. Dominant TE families were identical across CG and CHG DMTEs. 34.7% of CHH DMRs were TE-associated and were predominantly (86.2%) associated with hypermethylation. The most abundant CHH DMTE superfamilies were DTX (*unknown TIR*), XXX (unknown), DTT (*Mariner*), and DTH (*Harbinger*).

31 TE families were enriched among DMRs of at least one context (**Figure 4A**). TE families are referred to by their *clariTeRep* identifier and legacy ID (in brackets) where possible. Families enriched in more than one context included members of the *LTR Copia* superfamily: *RLC_famc15* (*HORPIA*), enriched among CG (4.5x) and CHG (3.3x) DMRs, and *RLC_famc16*, also enriched among CG (7.1x) and CHG (2.7x) DMRs. Both were primarily hypomethylated. Enriched Ty3-like families included *RLG_famc17*, enriched among CG (2.8x) and CHG (2.0x) DMRs, and *RLG_famc20*, enriched among CG (4.4x) and CHG (2.5x) DMRs; both tended towards hypomethylation in CG but hypermethylation in CHG contexts. Of unknown LTRs, *RLX_famc9* was enriched among CG (4.3x), CHG (1.9x), and CHH (25.4x) contexts, while *RLX_famc3* was enriched in CHG (1.7x) and CHH (4.4x). Both were predominantly hypermethylated in non-CG contexts, especially in CHH, while *RLX_famc9* was slightly associated with hypomethylated CG DMRs. *RLX_famc9* was the only TE family to be enriched among all contexts.

**Figure 4:**
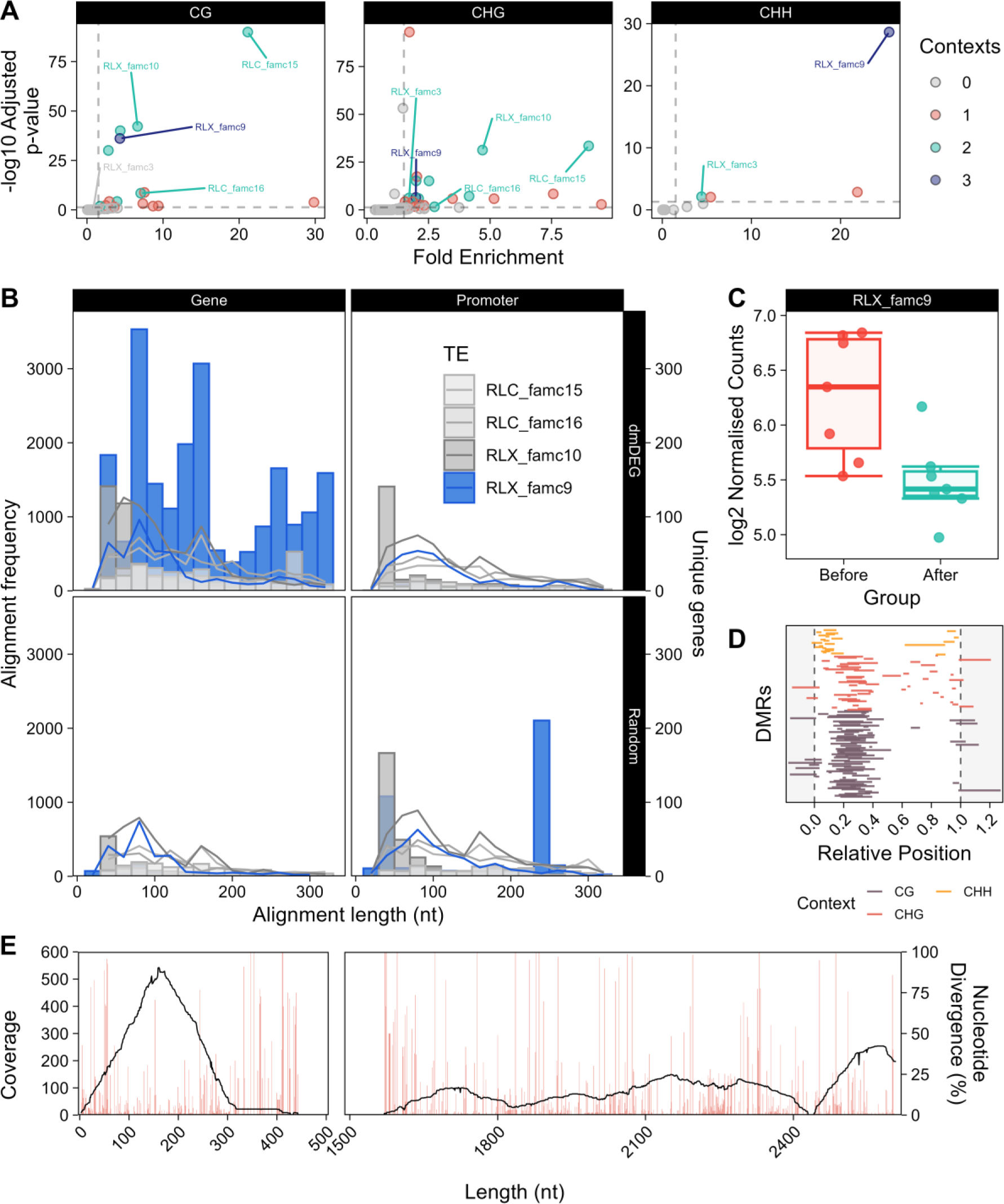
**(A)** DMR TE enrichment across all contexts showed several TE families to be enriched. Some TE families have been highlighted: families are coloured by their number of enriched DMR contexts (grey = 0, red = 1, green = 2, and blue = 3). **(B)** TE families enriched among DMRs were analysed for sequence similarity to the gene and promoter sequences of drought-associated genes (both methylome and transcriptome) and to random non-associated genes as a control. *RLX_famc9* was chosen due to its enrichment in all 3 C contexts, while *RLC_famc15*, *RLC_famc16*, and *RLX_famc10* were chosen due to their enrichment in 2 C contexts. Sequence similarity was plotted as a histogram of alignment lengths (left axis) and unique gene counts (right axis). **(C)** *RLX_famc9* transcript expression was compared before and after drought treatment. **(D)** Relative positions of DMRs overlapping with members of the *RLX_famc9* family were used to identify locations targeted for differential methylation. Relative positions were calculated for each DMR against the TE it was associated with. 0 represents the start of the TE; 1 represents the end. **(E)** Reads associated with *RLX_famc9* were mapped to a selected reference sequence; coverage (line chart, left axis) and percentage nucleotide divergence from the reference sequence (bar chart, right axis) were plotted against nucleotide position.

*RIX_famc7* (*Nadine*) and *RIX_famc9* (*Ramona*), both LINE transposons, were enriched among CG (4.0x and 1.7x respectively) and CHG (4.1x and 2.1x) DMRs; similarly to the Ty3-likes, they displayed substantial overlaps with hypomethylated DMRs.

Further enrichment results can be found in **Supplemental Data S4**.

### RLX_famc9 TEs show sequence similarity to differentially methylated and drought responsive genes

BLAST+ was used to identify sequence similarity between enriched TE families and the 1,290 promoters/ gene bodies of differentially methylated and differentially expressed genes. A random selection of 1,290 other genes and their promoters were used as a control set. Kolmogorov-Smirnov tests between alignment distributions at each nucleotide length (**Figure 4B**) identified *RLX_famc9* (*p = 1.06e-153*), *RLX_famc3* (*p = 4.85e-88*), and *RLC_famc16* (*p = 9.94e-56*) as the 3 most significant families for gene body alignments.

*RLX_famc9* displayed significantly higher homology to the target genes than the random genes, and was enriched in DMRs across all contexts. 63.2% of associated CG DMRs were hypomethylated, while 63.1% of non-CG DMRs were hypermethylated. GO terms including ‘*chloroplast stroma’*, ‘*chloroplast thylakoid membrane’*, ‘*starch biosynthetic process’*, and ‘*response to absence of light’* were significantly enriched in genes displaying gene body similarity to *RLX_famc9*; 55% of these were associated with hypomethylated CG DMRs.

34 differentially expressed TEs were identified. *RLX_famc9* transcripts were significantly downregulated (log₂FC = -0.85, *FDR = 0.044*) (**Figure 4C**). *RLX_famc9* DMRs were concentrated within the first third of their sequences (**Figure 4D**); CHH DMRs predominantly mapped in the first ∼10% of each sequence. Relative DMR position to the TEs was used to compare across the family. Alignments from all RNA samples to a representative *RLX_famc9* sequence were used to map transcriptionally active TE regions, identifying 2 detectable regions: a higher coverage region, assigned T1 (∼0-400 nt) and a lower coverage region, T2 (∼1600-2600 nt) (**Figure 4E**). Across all 1412 sites with associated reads, mean coverage was 128x, while mean divergence from the TE reference base was 7.94%.

TE transcripts may play a role in wheat’s drought response by producing sRNAs. We examined endoribonuclease *Dicer* and *Argonaute*-encoding gene expression (**Supplemental Data S5**) involved in cleavage and post-transcriptional gene silencing. Of 26 *Dicer* genes, 3 were upregulated after drought (*DCL4-B, DCL4-D*, and *RTL3*-like), while 3 (*DCL3a, DCL1-B*, and *DCL1-D*) were downregulated. Of 69 *Argonaute* genes, 4 were upregulated (*AGO2-A, AGO2-D, AGO18*, and *AGO4*) after drought and 1 was downregulated (*AGO-PNH1*).

## Discussion

In wheat, the physiological and transcriptomic impacts of drought are well understood, but the methylome impacts are undercharacterised. DNA methylation likely influences gene expression under biotic stress conditions ^28^, but its role in abiotic stresses like drought has been overlooked. Pooling genomic DNA for WGBS is an underutilised technique, despite its advantages for analysing complex genomes; usage of pooling has been carried out on small plant and animal genomes ^61–63^ but not used with genomes as large as wheat’s.

This effective and economical approach identified links between DNA methylation and gene expression shifts under drought, elucidating how the wheat drought response is regulated, and represents a step towards identifying epigenetic targets for crop improvement.

### TaROS1a homoeologues may contribute differently to the wheat drought response

*ROS1a* emerged as the most drought-responsive demethylases, consistent with reports that *ROS1* and *DME* genes are positively associated with drought tolerance in other species ^64,65^. Substantial CG/ CHG demethylation suggests active demethylation may play an important role in wheat’s drought response – something perhaps mediated by ROS1a, as suggested by the single homoeologue knockout TILLING lines for *ROS1a-5A, - 5B*, and *-5D*, potentially in a homoeologue-dependent way.

*ROS1a-5A* mutants appeared to exhibit significantly higher nDW than the wild-type allele-containing lines, while -5B and -5D mutants trended lower – though these comparisons were not significant – and all lines were more drought tolerant than WT Cadenza. The differences between our lines and Cadenza are likely due to the non-target mutation load present from the EMS, therefore we have primarily compared our *ROS1a* homozygous mutant lines to their homozygous wild-type out-segregants.

These contrasting results may suggest that the *ROS1a* homoeologues play distinct roles in mediating methylation under drought; *ROS1a-5A* may preferentially demethylate regions involved in growth restraint under water deficit, with its knockout here relieving this repression, while the -5B and -5D homoeologues may demethylate loci mobilised under drought, resulting in reduced drought tolerance when disrupted. Overall impacts across all three were modest, with a 1.7% reduction in nDW, though this could be due to compensatory homoeologue behaviour in polyploids ^66^, reducing the impact of detrimental mutations – though recent research has suggested that this assumption may not hold true in wheat ^67^. Single homoeologue knockouts for high level processes like epigenetic modulation may also have limited impacts.

*ROS1a* may contribute to drought tolerance in a homoeologue-specific manner, with *ROS1a-5A* disruption appearing to increase tolerance and disruption of its homoeologues associated with reduced nDW. Advanced backcrosses, double and triple knockouts of *ROS1a* homoeologues are needed to clarify their role in wheat’s drought response.

### Drought prompts massive transcriptional reprogramming accompanied by subtle epigenetic reprogramming of genic regions

Wheat’s transcriptomic response to drought has been substantially profiled ^21,68,69^; here we identified ∼14,000 DEGs (∼20% of expressed genes) between the BD and AD groups, indicating massive transcriptional reprogramming. By contrast, only ∼9% of genes were associated with DMRs, with the majority of regions associated with TEs and intergenic regions, especially among non-CG contexts, a response also seen in the rice methylome under drought ^70^. The modest genic response was not unexpected given many regulatory layers of the drought response. Hypermethylation was dominant in non-CG contexts; previous studies have identified hypermethylation in wheat leaves following drought events ^71,72^, though CG sites were substantially hypomethylated, potentially suggesting complex interplay or divergent roles across contexts.

Average methylation (CG ∼*88%*, CHG ∼*55%*, CHH *1.7%*) was congruent with previous estimates in wheat ^28^, while genic and TE methylation patterns were consistent with those seen in other species ^73,74^. We affirmed that genic CG methylation is significantly associated with gene expression: high gene body methylation is broadly a strong predictor of high gene expression, while high TSS methylation is a significant though weaker predictor of low gene expression. These findings are somewhat consistent with the concept of gene body methylation promoting maintaining high, stable expression ^25,75^ and high TSS/ first exon methylation inhibiting transcription ^76^.

Interestingly, genes with high log₂FCs were associated with substantially lower methylation at the TSS/ gene body, suggesting reduced DNA methylation may enable greater access by other transcriptional regulators under stress. Stress-responsive genes may exhibit substantial transcriptional noise in some organisms ^77^, something CG gene body methylation is believed to buffer in plants ^78^; we posit that low gene body methylation may be a characteristic of rapid stress response genes. Methylation could inhibit transcription factor target sequence binding accessibility, or promote histone H2A.Z exclusion ^79^, ultimately shielding a gene’s expression from environmental cues.

CG and non-CG DMRs displayed contrasting trends across the genome. Despite limited co-localisation, proximal DMRs across contexts were often positively correlated – especially CG and CHG DMRs – implying coordinated methylation control at those loci. CG and non-CG DMRs showed distinct methylation trends, primarily localised to different regions, and exhibited different traits, supporting a model in which each methylation context is governed by distinct mechanisms with distinct roles in controlling the genome.

### Genic regions undergo methylation changes in response to stress

Stress-associated functions were enriched in gene-body DMGs, suggesting that gene bodies are an RdDM or demethylation target under stress. Enriched GO terms describing signal transduction, phosphorylation, and hormone regulation were common across both hypo- and hypermethylated gene sets, indicating methylation may modulate these high-level processes.

Hypermethylated DMGs included many leucine-rich repeat domain-containing proteins, all involved in signalling and phytohormone responses. These were comprised of nucleotide-binding site leucine-rich repeat (*NBS-LRR*) disease resistance proteins, and leucine-rich repeat receptor-like kinases (*LRR-RLKs*). These *NBS-LRR*s included members of the *RGA* and *RPM* families, proteins which are implicated in stress-resistance pathways, including the gibberellic, salicylic, and abscisic acid cascades ^80–82^. Classically thought as pathogen responders ^83^, their stomatal control may make them useful methylation targets under water deficit. Among the hypermethylated *NBS-LRRs*, *RGA5* genes showed weak but broad upregulation across the family, suggesting they may be a methylation target. Similarly, *LRR-RLKs, including HPCA1s,* were upregulated; *HPCA1* encodes a plasma membrane-bound *LRR-RLK* implicated in ROS sensing and response within cells ^84^ and stress acclimation ^85^, while other identified *LRR-RLKs* were strongly associated with stress and growth mediation ^86^ and stress-responsive methylation changes ^87^.

Among upregulated hypermethylation-associated DMGs, triplet homoeologues of AAA-ATPases were identified, annotated with the enriched GO term ‘response to cold’, potentially suggesting sequence motif targeting by the RdDM pathway; AAA-ATPases are thought to play a role in salt and drought responses in maize, promoting drought resilience by inhibiting ROS accumulation ^88,89^.

Gene-flanking DMRs were enriched in TF binding motifs, indicating methylation may mediate expression by modifying TF binding. Methylation at binding sites can influence the TF binding affinity ^90,91^ – some TFs themselves are methylation-sensitive ^55,92^. Methyl groups may alter binding affinity by modulating steric hinderance or the minor groove width of DNA ^93^. Despite its potential regulatory role, methylation targeting TF motifs remains poorly studied.

Many stress-associated TFBMs were enriched among both gene-flanking hypo- and hypermethylated DMRs, perhaps due to motif variants within these large families being targeted by different RdDM mechanisms, or due to motifs varying in cytosine content. In *A. thaliana*, up to >75% of TFs are thought to be methylation-sensitive in some way ^94^, with methylation largely thought to inhibit TF binding. Several families thought to be methylation sensitive in *A. thaliana* were present here: bHLH and bZIP motifs were found to be strongly enriched among CHH promoter DMRs, suggesting that there may be CHH-mediated binding repression at these key stress-responsive families, while MYB and ERF/DREB TF motifs were enriched among DMRs across all contexts, suggesting that methylation may fine-tune the expression of specific gene responses. MYBs and their motifs are known to be regulated in part by methylation ^95^, with methylation playing a role in the recognition of their target sequences; ERF/DREB/AP2-associated motifs are sensitive to inhibition from hypermethylation ^94^, perhaps due to the high GC content of the DREB/CRT (GCCGAC) and GCC motifs (AGCCGCC) ^96^ associated with the superfamily.

Despite notable genes, most gene body DMGs showed little change in expression, supporting the idea that methylation is a subtle regulator at the genic level, stabilising high expression or potentially priming genes for future changes in expression by modulating the ability of other mechanisms to access a gene’s regulatory regions, as opposed to directly influencing expression. Sustained hypomethylation marks could prime genes for stress responsiveness ^97^, while gene body hypermethylation could shield genes from future environmental cues. We found no evidence of drought inducing methylation changes in promoters at the functional level, suggesting that promoter DMRs were not a widespread tool in wheat’s drought response, and that promoter-methylation-regulated genes would be better identified using a candidate-first approach.

### Specific transposable element families appear targeted for methylation changes

DMRs were found to be enriched in specific TE families. The role of TEs and TE-derived transcripts is understudied in wheat’s drought response; TE families in *A. thaliana* have been implicated in both *cis*- and *trans*-regulatory stress mechanisms ^31,98,99^, but similar epigenetically controlled responses in grasses remain elusive.

Enriched families such as *RLC_famc15, RIX_famc9*, and *RIX_famc7* are overrepresented in promoter sequences ^100^, though they were rarely accompanied by differences in the expression of proximal genes, suggesting that they did not play a large *cis*-regulatory role. Almost all genes in wheat’s genome have proximal TEs due to its high TE content, but previous studies have not identified strong associations between TE families and gene expression *cis*-regulation ^100^, suggesting that DMR-targeted TEs could instead possess *trans*-regulatory functions. It has been proposed that intron-localised TEs in *A. thaliana* and *Solanum lycopersicum* are associated with mostly enhanced gene expression of stress-associated genes ^101^, though we identified little association between TE methylation and associated gene expression.

### Methylation may mediate *RLX_famc9*-derived siRNAs

Some transposable elements are thought to regulate gene expression in *trans*. Methylation-mediated *trans*-acting TEs could act by modulating TF expression, indirectly influencing the expression of downstream genes ^102^, altering chromatin accessibility, potentially reducing the genic accessibility to proteins ^103^, or through the production of siRNAs from TE-derived mRNAs ^104,105^. We hypothesised that some wheat TEs, like *RLX_famc9*, can produce mRNA transcripts that are processed by the Post Transcriptional Gene Silencing (PTGS) pathway: DICER (DCL) endonucleases generate 21 or 24 nt siRNAs through cleavage, which are then loaded onto AGO-RISC complexes, and delivered to complementary transcripts for degradation, or to recruit DNA methyltransferases for the deposition of methylation at complementary loci ^106^.

Despite this, PTGS-associated DEGs indicated a reconfiguration of siRNA machinery, shifting towards *DCL4*/ *AGO2*-mediated PTGS through mRNA transcript disruption and away from *DCL3*-mediated RdDM silencing through methylation. Under drought, 21 nt sRNA-related genes (*AGO2*, *AGO18*, and *DCL4)* were upregulated. *AGO2s* load 21 nt siRNAs ^107^, while *AGO18* loads 21 nt phasiRNAs and some miRNAs ^108^, and *DCL4* is involved in the production of 21 nt siRNAs for mRNA transcript disruption ^109^. Conversely *DCL1s*, responsible for producing 21 nt miRNAs ^110^, were downregulated, suggesting increased siRNA and decreased miRNA production.

*DCL3*, the main *DCL* involved in cleaving TE transcripts to produce 24 nt siRNAs for RdDM ^111^, was also downregulated. A drought-induced shift from 24 nt siRNA-mediated RdDM, conventionally a major mechanism of TE methylation maintenance, towards rapid 21 nt-mediated PTGS may be favoured, as faster accumulation of mRNA-targeted siRNAs ^112^ could be advantageous for silencing detrimental stress-induced transcripts. A resultant decline in cellular 24 nt abundance would partially explain the increase in CG hypomethylation as methylation maintenance is scaled down.

*RLX_famc9* was the most promising potentially *trans*-acting TE family. RLXs are characterised by their long terminal repeat retrotransposon structure, but unlike the major RLC or RLG superfamilies, lack canonical gag or pol protein domains, and are considered ancient and more degenerated than other LTRs ^100^. Their lack of protein domain suggests an inability to transpose ^100^, which accompanied by their age (∼1.6 million years old), may have given rise to exaptation as stable functional regulatory elements within the genome ^113,114^. *RLX_famc9* showed significantly higher homology to drought-responsive genes when compared to random genes, indicating that it may play an important role in their regulation.

Reads and DMRs mapping to *RLX_famc9* sequences gave support for the presence of 1-2 methylation-regulated transcripts – reads mapped primarily to the TE’s 5’ end, while the relative positions of DMRs were densest within a similar range across the TE family, supporting methylation-regulation of a potential transcript. The T1 sequence was well conserved among the reads, suggesting that this region is present and stable across several members of the family. Though *RLX_famc9*’s associated CG and non-CG DMRs were opposing, greater hypermethylation in non-CG contexts at T1 may explain *RLX_famc9*’s downregulation; non-CG methylation – especially CHG – has been suggested as a stronger silencer of TEs than CG ^115^ and is critical for TE silencing ^116^. The potential reduction in 24 nt siRNAs could result in a reduction of TE self-silencing, which here could explain some of the hypomethylation occurring at CG sites ^117^, which are also preferentially targeted by the upregulated *ROS1a* glycosylases. Members of a TE family are considered highly homologous ^100,118^, suggesting that many individual TEs could be involved in transcripts generation; not all members of the TE family were subject to methylation changes, consistent with degradation. Additionally, the downregulation of *RLX_famc9*-derived transcripts, which we propose normally induce methylation at homologous drought-responsive genes, would contribute to their hypomethylation.

We propose that drought-inducible non-CG hypermethylation represses *RLX_famc9* transcription, reducing TE-derived siRNA production and abundance, and ultimately reducing RdDM activity at target genes, promoting hypomethylation as *de novo* methylation and maintenance rates slow. Coupled with ROS1a DNA glycosylase upregulation, this mechanism could enable greater drought-responsive gene expression modulation (**Figure 5**).

**Figure 5:**
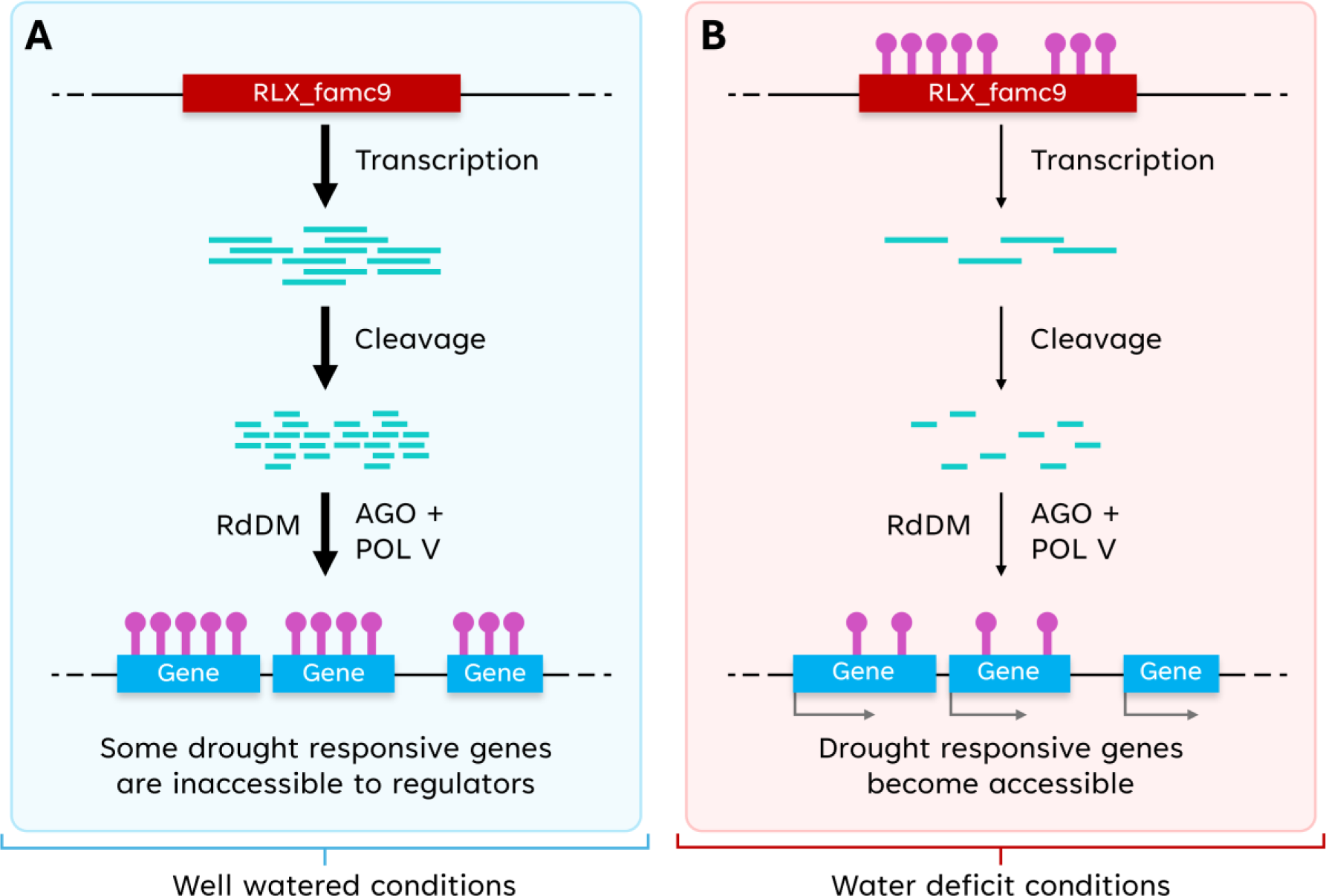
Our proposed mechanism of *RLX_famc9* action. Under well-watered conditions **(A)**, *RLX_famc9*-derived transcripts are freely produced due to hypomethylation of the transcriptionally active region. These dsRNA transcripts are cleaved into siRNAs by DICER proteins and loaded onto *AGO*/*Pol-V* complexes. These complexes recruit methyltransferases and are guided by the siRNA to deposit methylation at complementary loci, ultimately reducing their accessibility to transcriptional regulators. Under water-deficit conditions **(B)**, *RLX_famc9* TEs are hypermethylated to repress the expression of their transcriptionally active regions, reducing the transcription of dsRNAs and the abundance of 24 nt siRNAs. This allows for passive or active demethylation at the complementary genes, increasing their accessibility to transcriptional regulators in response to drought. *RLX_famc9*-derived transcripts are represented in blue, while methylated loci are represented in purple.

Our findings highlight the multifaceted role of DNA methylation in wheat’s drought response. Gene body methylation appears to support stable, stress-insensitive expression, while promoter methylation appears to have limited impact, implying that methylation-mediated gene control is more targeted and complex than previously assumed. Transcription factor binding motifs enriched in DMRs pointed towards a mechanism in which methylation modulates binding affinity, especially among genes regulated by the MYB, bZIP, bHLH, and ERF/DREB TF families. The vast majority of differential methylation, however, occurred in intergenic space: transposons emerged as candidates for *trans*-acting regulation. We propose a model in which the *RLX_famc9* TE, which was differentially methylated and differentially expressed under drought, and displayed significant sequence similarity to drought responsive genes, may play a repressive *trans-*regulatory role under normal conditions, but be silenced under drought to relieve this repression.

Our analysis also demonstrates that pooled DNA methylation sequencing in complex polyploids like wheat is effective, yielding high-quality data and clear, functional methylation divergences between samples at reduced cost. Future work with more samples, accompanied by siRNAs profiling, will be crucial for exploring our proposed mechanisms and testing *RLX_famc9*’s proposed stress-associated gene regulation.

## Materials and Methods

### Plant growth

The details of the plant growth conditions were reported previously in Barratt & Reynolds et. al (2023), though only half of the landraces (7 lines) from that study were selected for further investigation in this study. YoGI landrace lines deemed to exhibit a drought tolerant phenotype (YoGI_007, YoGI_021, YoGI_026, YoGI_047, YoGI_059, YoGI_145, and YoGI_261), defined as maintaining >70% biomass by weight compared to their control group, were selected from the wider panel of 20 as they were hypothesised to more likely exhibit beneficial drought-response mechanisms than susceptible lines. Four biological replicates were grown per line.

In summary, plants were sown in an 80:20 Levington’s F2/ sand mix treated with CaLypso insecticide (Bayer CropScience Ltd., 0.083ml in 100ml water, applied per litre of compost), grown in long-day conditions (16/8h) and commenced drought conditions upon reaching the growth stage GS13 (Zadok’s growth scale; ^34^). Drought treatment consisted of withholding water for 10 days, followed by 3 days of recovery with regular watering. Plants were sampled for RNA and DNA analyses at onset of drought (day 0) and end of drought (day 10). Dry biomass of above-ground shoots from droughted samples was compared to the above-ground biomass from control samples. Soil moisture content was recorded using an ML3 Thetaprobe Soil Moisture Sensor with an HH2 Moisture Meter (Delta-T Devices, Cambridge, United Kingdom) at onset and end of treatment, as well as at the end of recovery.

### TILLING Mutant acquisition, backcrossing, and drought screening

TILLING mutant lines for *TaROS1a* homoeologues were obtained from SeedStor (https://seedstor.ac.uk/). Identification of individual lines with mutations in the genes of interest (*ROS1a-5A*: *TraesCS5A02G169000*; *ROS1a-5B: TraesCS5B02G165800*; *ROS1a-5D: TraesCS5D02G173300*) was carried out using the EnsemblPlants variant tables (https://plants.ensembl.org/Triticum_aestivum) available for each gene. These genes have previously been identified as members of the *TaROS1a* family ^35^ but have also been referred to as *DME* orthologues. Variants with a Cadenza background were selected based on the presence of a predicted nonsense mutation (gain of stop codon). If multiple nonsense mutant lines were identified, the one with the mutation earlier in the sequence was selected to reduce the chance of partially functional but truncated transcripts.

*Cadenza0002, Cadenza1580*, and *Cadenza1622* were selected due to the presence of nonsense mutations for *ROS1a-5A, -5B*, and -*5D* respectively. These lines were originally generated using EMS mutagenesis in the mid-2010s ^36^ but current seed stocks may be a result of several generations of self-pollination. TILLING mutants were confirmed using polymerase chain reaction (PCR)-based genotyping (**Supplemental Data S1**) and sequencing. Polyploids often contain multiple homoeologues of a given gene across its genomes; homoeologue-specific primers are needed to amplify only the region of interest on the gene of interest. Primers were designed using Primer3 ^37^ and screened for GC content, appropriate melting temperature, and the likelihood of self-dimerisation or hairpin formation. Primers were also tested for specificity against the homoeologues using NCBI’s primer-BLAST tool: non-homoeologue specific primers were noted and the nucleotide at the site of interest was checked to ensure that homoeologues could be differentiated when examining the sequencing trace at the genotyping stage. Primers were reconstituted as per the manufacturer’s instructions, and a working aliquot was made at 10 µM concentration. Forward and reverse primer sequences are available in **Supplemental Data S1**.

PCR was carried out on DNA extracted from leaf tissue (harvested, immersed in liquid nitrogen, and stored at -80°c) using an Edwards DNA Extraction buffer. GoTaq Green Master Mix (MM) (Promega, WI, USA) was used for PCR amplification. Each PCR reaction contained 10 µL GoTaq MM, 1 µL of DNA, 1 µL Forward Primer, 1 µL Reverse Primer, and 7 µL H_2_O to a total volume of 20 µL. PCRs were carried out using the same cycle quantity and length, though primer annealing temperatures differed due to GC proportions. PCR products were purified prior to sequencing using the Wizard SV Gel and PCR Cleanup Kit (Promega, WI, USA) as per the manufacturer’s protocols. 2.5 µL of purified PCR product was added to 5 µL of nuclease-free H_2_O with 2.5 µL of Forward Primer to a total volume of 10 µL and was sent for sequencing using Eurofins’ LightRun Tube Sanger Sequencing service. Presence of mutation at the desired site was confirmed and plants were designated ROS1a-5X^WT^ (out-segregants; no point mutation) or ROS1a-5X^MUT^ (point mutation present).

Confirmed mutants were backcrossed onto wild-type Cadenza plants. Mutant lines were used as pollen donors. Backcrossing was carried out to reduce the mutation load of the progeny while retaining the mutation of interest. Experimental plants were backcrossed once, and the progeny was self-pollinated to obtain homozygous mutants for each line. BC_1_S_1_ (once backcrossed, once selfed) plants were used for drought screening. Plants were genotyped at each stage to ensure that the lines maintained the mutation of interest as described previously.

BC_1_S_1_ and WT Cadenza plants were grown and screened for drought tolerance as previously described. Cadenza was used to identify a baseline WT drought tolerance prior to EMS mutagenesis and backcrossing. Mutant BC_1_S_1_ lines were compared to their WT-allele siblings and WT Cadenza using normalised dry weight (*nDW*) as a measure of drought tolerance across the different experimental lines. *nDW* was calculated as 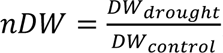, where *DW*_*drought*_ refers to an individual plant’s dry weight under drought conditions and *DW*_*drought*_ represents the mean dry weight for that line under control conditions. *nDW* is expressed as a proportion.

Statistical testing was carried out compare traits between the ROS1a-5X^WT^, ROS1a-5X^MUT^, and WT Cadenza lines. Data were assessed for homogeneity of variance and normality of residuals using the shapiro_test() and levene_test() functions from the rstatix (ver. 0.7.2) ^38^ package in R. If assumptions were fulfilled, a two-way Analysis of Variance (ANOVA) test was used in R followed by the TukeyHSD() function; if assumptions were violated, the non-parametric Aligned Rank Transformed ANOVA (ver. 0.11.2) (ART-ANOVA) from the ARTool ^39^ package was used, followed by post-hoc Tukey testing via the ARTool art.con() function.

### Nucleic acid extraction and sequencing

The details of the RNA extraction protocols have been reported previously in ^21^. In summary, RNA was extracted from ∼100 mg of individual leaf tissue before and after drought and sequenced on the Illumina Novaseq 6000 platform (Illumina, CA, USA).

Genomic DNA was extracted using the Wizard HMW DNA Extraction Kit (Promega, WI, USA) according to the manufacturer’s protocol, including an RNAse treatment. Genomic DNA concentration was quantified using a Qubit dsDNA Broad Range Assay Kit and Qubit 4 Fluorometer (Life Technologies, CA, USA), with DNA quality and integrity quantified using a NanoDrop ND-1000 Spectrophotometer (Thermo-Fisher Scientific, MA, USA) and an Agilent Technology Tapestation (Agilent Technologies, CA, USA); samples were only included where their DNA integrity score (DIN) was above 7. DNA samples were pooled together in equimolar proportions into two pools, before drought (BD) and after drought (AD). Samples were then shipped on dry ice to Novogene (Cambridge, United Kingdom) for further QC, bisulphite treatment, library preparation, and sequencing on the Illumina Novaseq 6000 platform (150 bp paired end strategy).

### Genomic data processing, mapping, and QC

Post-sequencing quality control was carried out using FastQC (version 0.11.9; www.bioinformatics.babraham.ac.uk/projects/fastqc/) for both RNA and genomic DNA reads. Raw reads were filtered using Trim Galore ^40^ (version 0.6.10) by trimming low quality sequences (average Phred score < 20), trimming short length reads (<20bp) and clipping Illumina adapters. For DNA, 8 bp from the 5’ end of each read was trimmed to avoid sequencing/ methylation biases. FastQC was also run post-trimming to verify adapter and poor-quality sequence removal.

Further RNA processing was previously detailed in ^21^; in summary, reads were aligned using Salmon ^41^ to the IWGSC *Triticum aestivum* v1.1 transcriptome reference.

Genomic DNA sequence processing, including alignment and methylation calling, was carried out using Bismark ^42,43^. The genome assembly was pre-processed using the *bismark_genome_preparation* script at default parameters. Paired-end Bismark alignment (bowtie2) was run at mostly default settings (*parameters: ‘--score_min L,0,-0.4’*) with slightly relaxed penalties to account for differences between the landraces and Chinese Spring. Reads were mapped to the IWGSC *Triticum aestivum* v1.0 assembly (GCA Accession: GCA_900519105.1); the IWGSC *Triticum aestivum* v1.1 annotation was used for annotating features. Trimmed sequences were aligned using end-to-end alignment mode. Following alignment, the Bismark script *deduplicate_bismark* was used for deduplication of reads, after which *bismark2bedGraph* (*parameters: ‘--CX’, ‘--ample_memory’*) was used for methylation extraction for all three methylation contexts for both samples.

Coverage and sequencing depth of genomic reads was assessed using PanDepth ^44^ following alignment.

### Transcriptome analysis

The differential expression analysis protocol was previously detailed in ^21^; count data from Salmon was analysed using DESeq2 ^45^ after basic filtering of data. Differential expression analysis was carried out using BD samples as the reference level, and resultant log₂Fold Change values were shrunk using the R package Ashr ^46^. Thresholds of absolute log₂Fold Change ≥ 1 and FDR-adjusted p-value ≤ 0.05 were used to screen for differentially expressed genes (DEGs).

### BS-Seq, initial visualisation, and differential methylation analysis

After alignment, Bismark was used to generate Bismark Coverage files, which contained methylation percentage, unmethylated reads, and total reads for each sequenced site per chromosome in each sample.

Methylation extraction for all four contexts was run using default parameters apart from ‘*--ample_memory’* and ‘*--cutoff 3’*. A cutoff of 3 was used to filter out very-low coverage sites.

Methylation data across gene bodies, TE bodies, and their flanking regions was processed using the *MethOverRegion* function from the ViewBS package ^47^, and then refined and visualised using R. Custom scripts were used to visualise and analyse methylation traces across gene bodies per percentile of gene expression and absolute log₂Fold Change.

Differential methylation analysis (DMA) was performed using the R package DSS ^48^. DSS input was filtered to remove loci that with coverage less than 3 in one or both samples. The DMA was carried out using the DMLtest() function with smoothing at default parameters. As nearby cytosine methylation sites are assumed to be spatially correlated ^48,49^, smoothing was used to improve DML and DMR detection.

DMLs and DMLRs obtained using DSS were filtered using thresholds for each sequence context. For all comparisons, a p-value threshold of 0.05 was used. For CG and CHG methylation, DMLs were considered significant with absolute Δ methylation ≥ 20%, while the threshold for CHH DMLs was absolute Δ methylation ≥ 10%. Different thresholds were used due to the different frequency of each type of site and average methylation of each context across the genome; CHH sites are more numerous than CHG or CG sites but are methylated at significantly reduced levels. Thresholds for DMR filtering were identical, though DSS’s *areaStat* statistic was used for ad-hoc DMR-ranking instead of p-values as DSS does not provide p values for DMRs. AreaStat is a function of test statistics incorporating width (number of DMLs in the region) and height (Δ methylation). Very C-poor DMRs (less than 5% DMLs) were removed from downstream analysis. Genes overlapping with DMRs by at least 10bp within the gene body or 1.5 kb flanking intergenic regions were defined as differentially methylated genes (DMGs). Transposable elements that overlapped with DMRs by at least 10bp were defined as differentially methylated transposable elements (DMTEs). Overlapping elements were identified using genomation ^50^ and valr ^51^ in R.

### Gene ontology and transcription factor binding motif enrichment analysis

Gene Ontology enrichment analysis (GO) of DEGs and DMGs was carried out using clusterProfiler (v 4.12.0) ^52^ against the relevant background for each data set (DEGs against all genes remaining post filtering; DMGs against all wheat genes with GO terms and coverage). GO terms with an FDR-adjusted *p* value ≤ 0.05 by hypergeometric testing were considered significantly enriched.

Transcription Factor Binding Motif enrichment analysis was conducted using the findMotifsGenome.pl script from HOMER2 ^53^. BED files for promoter and downstream DMRs, as well as the genome reference, were supplied as input. Parameters included ‘-*modelBg’* and ‘-*mset’ plants.* For the -size argument, the average length of the flanking DMRs was given for each context (CG: ‘-*size -278,278’*; CHG: ‘-*size -262,262’*; CHH: ‘-*size -171,171’*) to include motifs both nearby and overlapped by the DMRs; methylation levels surrounding TF binding sites is thought to be associated with TF binding affinity ^54,55^.

### Transposable enrichment and homology analysis

Enrichment analysis of differential methylation in transposable element families was carried out using the TEENA pipeline ^56^, using all DMRs obtained per context from the differential methylation analysis. All DMRs were used to identify TE families selected for any sort of differential methylation activity, as TE families were unlikely to be differentially methylated in one direction only. The TE annotation was summarised to the family level, grouping together all subfamilies due to their high homology. TE families were considered significantly enriched if they had an adjusted p-value ≤ 0.05 and a fold enrichment ≥ 2.

BLAST+ ^57^ (v2.14.1) was then used to identify TE families with high homology to the 1.5 kb promoter regions and gene bodies of differently methylated and differentially expressed genes. Genomic sequences of the TEs were also queried against a random gene set of equal size as a control. Each gene set was converted into two BLAST *nucl* databases (one for promoter regions, one for gene bodies) using the function *makeblastdb* (parameters: ‘-parse_seqids’, ‘-dbtype nucl’). TE sequences were obtained from the Wheat RefSeq v1.1 annotation using BEDTools ^58^ and queried against the individual gene set databases using BLAST+ blastn (parameters ‘*-word_size 8’, ‘-evalue 0.0005’)* for both gene bodies and promoter regions. Results were filtered to remove alignments below 19 nt and above 320 nt, as well as removing genes/ promoter regions that overlapped with the queried transposable element family. Alignments were visualised using histograms in ggplot2 alongside the number of unique genes per histogram bin. TEs that showed significantly more alignment to the target genes than the random genes were identified using the function *ks.test*() to carry out a Kolmogorov-Smirnov test in R.

Orthologous TEs were found in the TRansposable Elements Platform (TREP) database to the TEs of interest from the IWGSC V1.1 reference annotation using BLAST+. RNA-seq reads were then mapped to the TREP Complete nucleotide sequence dataset ^59^ with gene CDS models supplied as decoys for alignment to ensure that reads preferentially mapped to TEs rather than genes. The TREP database contains unique and consensus transposable element sequences from across the *Triticeae*. Salmon was run with default parameters. Quantified expression was processed as previously described for gene expression data and analysed using DESeq2 ^45^.

A multiple sequence alignment of RNA reads and an *RLX_famc9* reference sequence was carried out using MAFFT ^60^ with the parameters ‘*mafft --reorder --keeplength –anysymbol --maxambiguous 0.05 --kimura 1 --addfragments fragments --auto’*. The *RLX_famc9* sequence was selected through all-vs-all comparison of the *RLX_famc9* family sequences using BLAST. After self-hit removal, bitscore was calculated for each sequence. The final sequence was selected by identifying the sequence closest to the mean sequence length within the 95^th^ or above percentile. The final sequence used is available in **Supplemental Data S6**.

## Supporting information

Supplementary data (all)

## Data availability

The datasets used in this study can be found in online repositories. Raw and processed data is publicly available at NCBI GEO (RNA-seq data: GSE225797; WGBS data: GSE311296).

The Cadenza TILLING lines used (*Cadenza0002*, *Cadenza1622*, and *Cadenza1580*) are available from the Germplasm Resources Unit (GRU) (Norwich, United Kingdom; www.seedstor.ac.uk).

Custom scripts used in this article are available at https://github.com/andreaharper/HarperLabScripts.

## Author contributions

ALH conceived and supervised the project. IJR and LJB designed and performed the experiments. IJR wrote the original draft of the manuscript. All authors reviewed, edited, and approved the final manuscript.

## Acknowledgements

The authors thank the John Innes Centre Germplasm Resources Unit, a National Bioscience Research Infrastructure supported by the UKRI-BBSRC (grant: BBS/E/JI/23NB0001) for conserving and supplying TILLING germplasm through www.seedstor.ac.uk, as well as CIMMYT, Mexico, and the Crop Research Institute, Czechia for providing additional germplasm.

